# High-content screening for rare respiratory diseases: readthrough therapy in primary ciliary dyskinesia

**DOI:** 10.1101/2020.02.28.959189

**Authors:** Dani Do Hyang Lee, Daniela Cardinale, Ersilia Nigro, Colin R. Butler, Andrew Rutman, Mahmoud R. Fassad, Robert A. Hirst, Dale Moulding, Alexander Agrotis, Elisabeth Forsythe, Daniel Peckham, Evie Robson, Claire M. Smith, Satyanarayana Somavarapu, Philip L. Beales, Stephen L. Hart, Sam M. Janes, Hannah M. Mitchison, Robin Ketteler, Robert E. Hynds, Christopher O’Callaghan

## Abstract

Development of therapeutic approaches for rare respiratory diseases is hampered by the lack of systems that allow medium-to-high-throughput screening of fully differentiated respiratory epithelium from affected patients. This is a particular problem for primary ciliary dyskinesia (PCD), a rare genetic disease caused by mutations in genes that adversely affect ciliary movement and consequently mucociliary transport. Primary cell culture of basal epithelial cells from nasal brush biopsies, followed by ciliated differentiation at air-liquid interface (ALI) has proven to be a useful tool in PCD diagnostics but the technique’s broader utility, including in pre-clinical PCD research, has been limited by the number of basal cells that it is possible to expand from such biopsies. Here, we describe a high-content, imaging-based screening method, enabled by extensive expansion of PCD patient basal cells and their culture into differentiated human respiratory epithelium in miniaturised 96-well transwell format ALI cultures. Analyses of ciliary beat pattern, beat frequency and ultrastructure indicate that a range of different PCD defects are retained in these cultures. We perform a proof-of-principle personalized investigation in reduced generation of motile cilia (RGMC), a rare and very severe form of PCD, in this case caused by a homozygous nonsense mutation (c.441C>A; p.Cys147*) in the *MCIDAS* gene. The screening system allowed multiple drugs inducing translational readthrough to be evaluated alone or in combination with inhibitors of nonsense-mediated decay. Restoration of basal body formation in the patient’s nasal epithelial cells was seen *in vitro*, suggesting a novel avenue for drug evaluation and development in PCD.

**Summary:** We describe primary cell culture of nasal epithelial cells from patients with primary ciliary dyskinesia including differentiatiation of these to a ciliary phenotype and high-content screening in miniaturised air-liquid interface cultures.

## Introduction

Primary ciliary dyskinesia (PCD) is a rare autosomal recessive genetic disorder arising from abnormalities in the structure and function of motile cilia. The disease is characterized by recurrent respiratory tract infections and early onset bronchiectasis, chronic nasal symptoms and sinusitis and in many a significant hearing deficit due to glue ear. Genetically, PCD is a heterogeneous disorder with causative variants in more than 40 different genes accounting for disease in around 70% of PCD patients [1–3]. While the specific ultrastructural defects caused by these mutations vary, they generally occur in genes that encode proteins involved in axonemal structure, regulatory complexes, ciliary assembly or ciliary transport [4]. Mutations frequently affect the outer dynein arm and cause ciliary stasis or a combination of ciliary stasis and marked reduction in ciliary beat amplitude, but a wide range of ciliary beat defects have been observed, including reduced amplitude, rotational beating and hyperkinesia.

As a result of disease diversity, there is no single PCD diagnostic test: clinical history, genetic analyses, low nasal nitric oxide levels and the direct observation by light microscopy of abnormal ciliated epithelial cells in nasal brush biopsies using high-speed video analysis as well as analyses of ciliary fine structure by electron microscopy are used, often in combination, to reach a PCD diagnosis [5]. In specialised centres, cell culture is used to improve the diagnosis because ciliary dyskinesia secondary to infection and inflammation can make interpretation difficult. Examining cultured cells can also reduce the requirement for re-biopsy when biopsy samples do not contain ciliated cells [6] and is advantageous when characterizing atypical PCD phenotypes such as inner dynein arm defects, ciliary disorientation or reduced generation of multiple motile cilia.

The multiplicity of patient genotypes and phenotypes in PCD suggest it as a fertile ground for the personalized medicine approach. However, nasal epithelial cell culture from brush biopsies of PCD patients has been limited by the use of growth media in which basal cells undergo senescence soon after culture initiation. As few cells are obtained during nasal or lower airway biopsy procedures, the majority of investigations use cells directly from air-liquid interface (ALI) cultures (where cells are cultured in transwells and fed basally in order to expose them to air) or after minimal passaging [7], largely precluding the biobanking of primary PCD cell cultures. Improvements in epithelial cell culture have led to methods that allow extensive expansion of airway [8] and nasal [9] epithelial cells from healthy donors and patients with genetic diseases such as cystic fibrosis [10–12]. In PCD, a previous study of *RSPH1*-mutant PCD patients indicated that patient phenotypes are recapitulated upon differentiation *in vitro* [13]. Here, we demonstrate extensive expansion of basal cells from PCD patients with diverse causative mutations in 3T3-J2 fibroblast feeder cell co-culture in medium containing Y-27632, a Rho-associated protein kinase (ROCK) inhibitor, and the maintenance of the patients’ ciliary phenotypes in ALI cultures. We also miniaturise ALI cultures from PCD patients and healthy volunteers to a 96-well format that allows high-content screening. Around a quarter of PCD patients carry nonsense mutations that introduce a premature termination codon (PTC) into the mRNA, leading to an absence of the protein or the production of inactive, truncated forms [14]. PTC-‘readthrough’ can bypass the PTC, leading to partial or full expression of functional protein [15]. Several readthrough agents have been described, including the antibiotic gentamycin and ataluren, a drug in development for Duchenne muscular dystrophy [16] and that has also been trialled in cystic fibrosis [17]. Analysis of PCD PTCs cloned into luciferase reporter constructs suggested that a subset of PTCs could be suppressed by aminoglycoside readthrough agents *in vitro* [18].

Here, we focus on one of our patients who has a reduced generation of multiple motile cilia (RGMC) ciliopathy, caused by a nonsense mutation in *MCIDAS* – the gene encoding multicillin, a transcriptional co-factor that regulates early steps in ciliogenesis [19]. The patient’s genetic screening and *MCIDAS* mutation (c.441C>A (p.Cys147*) have been reported previously [20]. We use a high-content immunofluorescence screening approach to assess the formation of multiple basal bodies – structures that organise microtubules during multi-ciliogenesis – in miniaturised ALI cultures. Despite the observation of isolated basal bodies in nasal brushing from patients with this condition, the formation of multiple basal bodies is compromised by the lack of this protein in ALI cultures [20, 21]. In a semi-automated ALI culture protocol, combinations of readthrough agents and inhibitors of nonsense-mediated decay (NMD) were able to restore formation of basal bodies – but not cilia – in the patient’s epithelial cells.

## Results

### Cell culture expansion of PCD patient basal epithelial cells

Brush biopsy-derived nasal epithelial cells were cultured from 13 PCD patients with a range of genetic defects and 11 healthy donors with no known ciliary defects (Table 2). After isolation in BEGM [22, 23], cultures from four healthy donors and ten PCD patients were expanded separately in both BEGM (Figure 1A) and in co-culture with 3T3-J2 mouse embryonic fibroblast feeder cells in the presence of a Rho-associated protein kinase (ROCK) inhibitor (Y-27632; 3T3+Y) (Figure 1B). In BEGM, nasal epithelial cells reached senescence before 10 population doublings in both healthy (Figure 1C) and PCD cultures (Figure 1D). In contrast, and consistent with previous observations [9], expansion in 3T3+Y was rapid and was sustained for at least 8 passages (Figure 1C, D). 3T3+Y cultures from healthy and PCD donors were morphologically indistinguishable and, as expected, PCD cultures expressed the basal cell-associated proteins keratin 5 and P63 (Figure 1E). Further, comparing the abundance of the basal cell-associated mRNAs *KRT5* (Figure 1F), *KRT14* (Figure 1G), *P63* (Figure 1H), *ITGA6* (Figure 1I) and *NGFR* (Figure 1J) between control and PCD nasal cultures confirmed that cells in PCD cultures are basal epithelial stem/progenitor cells.

**Figure 1:**
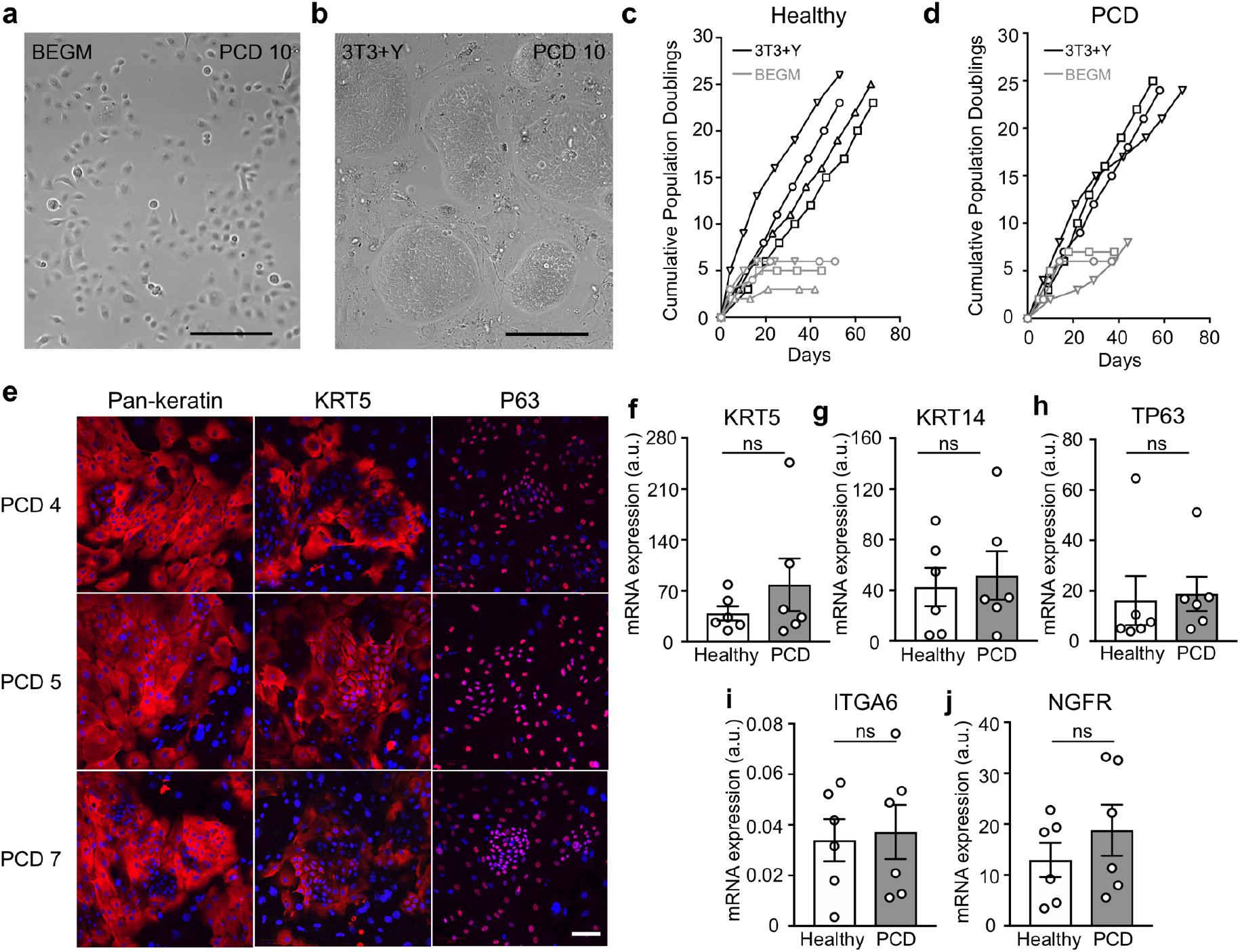
Expansion of nasal basal epithelial cells from patients with primary ciliary dyskinesia. (A) Brightfield microscopy showing a representative example of nasal epithelial cells in BEGM culture (PCD 10) at passage 1. (B) Representative example of brightfield microscopy showing nasal epithelial cells (PCD 10) in 3T3+Y culture at passage 2. Scale bar = 100µm. Part of this image previously appeared in a schematic diagram in a published methods chapter [24]. Reprinted by permission from Springer Nature: Springer eBook, Methods in Molecular Biology, Springer Science Business Media New York (2019). (C) Cumulative population doublings for four healthy donor nasal epithelial cell cultures in BEGM (grey) and 3T3+Y (black). (D) Cumulative population doublings for three primary ciliary dyskinesia (PCD) patient nasal epithelial cells in BEGM (grey) and 3T3+Y (black). (E) Representative examples of immunofluorescence images showing pan-keratin, keratin 5 and P63 expression (red) in three PCD patient 3T3+Y cultures. The nuclear counterstain DAPI is shown (blue). Scale bar = 100 µm. (F-J) qPCR analysis comparing the expression of keratin 5 (*KRT5*; F), *KRT14* (G), *TP63* (H), integrin alpha 6 (*ITGA6*; I) and nerve growth factor receptor (*NGFR*; J) between healthy and PCD donor basal cell cultures in 3T3+Y. *P* values were derived using two-tailed paired t-tests, ns indicates *P* > 0.05.

**Table 2:**
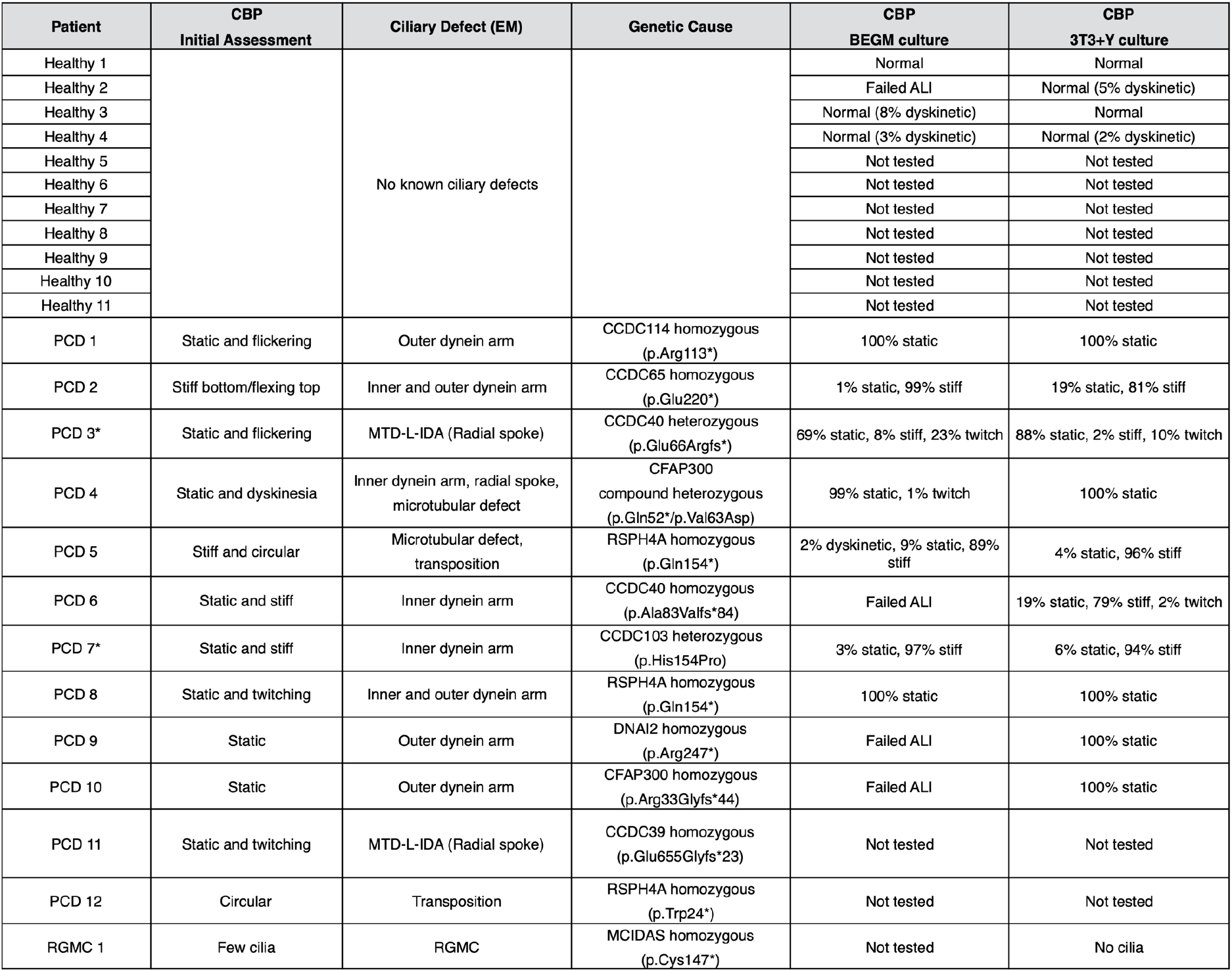
Subject-specific ciliary beat pattern in biopsies and cell cultures with primary ciliary dyskinesia. Asterisks indicate two cases where a single variant was found, but with a typical PCD diagnosis and ciliary defects that are recognised as consistent with being linked to the affected gene. MTD-L-IDA = microtubules disorganisation lack of internal dynein arm.

| Patient | CBP<br>Initial Assessment | Ciliary Defect (EM) | Genetic Cause | CBP<br>BEGM culture | CBP<br>3T3+Y culture |
| --- | --- | --- | --- | --- | --- |
| Healthy 1 |  | No known ciliary defects |  | Normal | Normal |
| Healthy 2 |  |  |  | Failed ALI | Normal (5% dyskinetic) |
| Healthy 3 |  |  |  | Normal (8% dyskinetic) | Normal |
| Healthy 4 |  |  |  | Normal (3% dyskinetic) | Normal (2% dyskinetic) |
| Healthy 5 |  |  |  | Not tested | Not tested |
| Healthy 6 |  |  |  | Not tested | Not tested |
| Healthy 7 |  |  |  | Not tested | Not tested |
| Healthy 8 |  |  |  | Not tested | Not tested |
| Healthy 9 |  |  |  | Not tested | Not tested |
| Healthy 10 |  |  |  | Not tested | Not tested |
| Healthy 11 |  |  |  | Not tested | Not tested |
| PCD 1 | Static and flickering | Outer dynein arm | CCDC114 homozygous<br>(p.Arg113*) | 100% static | 100% static |
| PCD 2 | Stiff bottom/flexing top | Inner and outer dynein arm | CCDC65 homozygous<br>(p.Glu220*) | 1% static, 99% stiff | 19% static, 81% stiff |
| PCD 3* | Static and flickering | MTD-L-IDA (Radial spoke) | CCDC40 heterozygous<br>(p.Glu66Argfs*) | 69% static, 8% stiff, 23% twitch | 88% static, 2% stiff, 10% twitch |
| PCD 4 | Static and dyskinesia | Inner dynein arm, radial spoke,<br>microtubular defect | CFAP300<br>compound heterozygous<br>(p.Gln52*/p.Val63Asp) | 99% static, 1% twitch | 100% static |
| PCD 5 | Stiff and circular | Microtubular defect,<br>transposition | RSPH4A homozygous<br>(p.Gln154*) | 2% dyskinetic, 9% static, 89%<br>stiff | 4% static, 96% stiff |
| PCD 6 | Static and stiff | Inner dynein arm | CCDC40 homozygous<br>(p.Ala83Valfs*84) | Failed ALI | 19% static, 79% stiff, 2% twitch |
| PCD 7* | Static and stiff | Inner dynein arm | CCDC103 heterozygous<br>(p.His154Pro) | 3% static, 97% stiff | 6% static, 94% stiff |
| PCD 8 | Static and twitching | Inner and outer dynein arm | RSPH4A homozygous<br>(p.Gln154*) | 100% static | 100% static |
| PCD 9 | Static | Outer dynein arm | DNAI2 homozygous<br>(p.Arg247*) | Failed ALI | 100% static |
| PCD 10 | Static | Outer dynein arm | CFAP300 homozygous<br>(p.Arg33Glyfs*44) | Failed ALI | 100% static |
| PCD 11 | Static and twitching | MTD-L-IDA (Radial spoke) | CCDC39 homozygous<br>(p.Glu655Glyfs*23) | Not tested | Not tested |
| PCD 12 | Circular | Transposition | RSPH4A homozygous<br>(p.Trp24*) | Not tested | Not tested |
| RGMC 1 | Few cilia | RGMC | MCIDAS homozygous<br>(p.Cys147*) | Not tested | No cilia |

### Co-cultured PCD basal cells form mucociliary epithelia at an air-liquid interface

Having established a method to extensively expand and cryopreserve nasal epithelial cells from PCD donors, we optimized methods to differentiate these at an air-liquid interface (ALI; Figure 2A). This method allowed us to assess their ciliary phenotypes [8, 20, 22, 24]. PCD cells derived from 3T3+Y cultures formed confluent ALI cultures comparably to healthy controls and the barrier function of healthy and PCD-derived ALI cultures were similar, as assessed by trans-epithelial electrical resistance (TEER) measurements taken once cultures were fully differentiated (Figure 2B). After 5 weeks of ALI culture, the expression of the basal cell-associated genes *KRT5* (Figure 2C) and *P63* (Figure 2D) had markedly decreased and there were trends towards increased expression of the differentiation-associated *KRT8* (Figure 2E), *MUC5AC* (Figure 2F), *MUC5B* (Figure 2G) and *FOXJ1* (a transcription factor expressed within the ciliated cell lineage; Figure 2H), consistent with the expected differentiation of basal cell cultures towards mature epithelia containing ciliated cells and mucosecretory cells.

**Figure 2:**
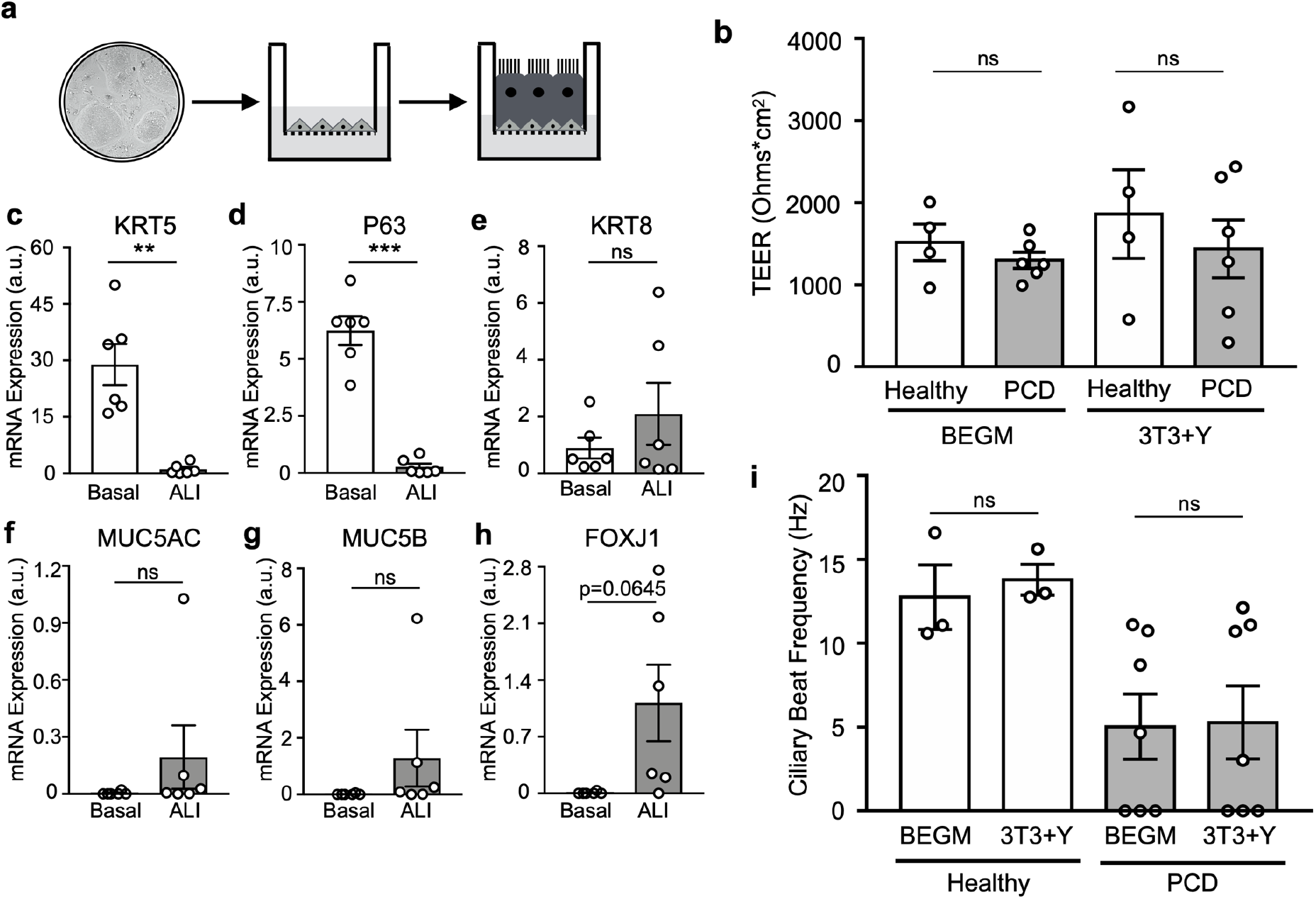
Ciliogenesis in cultured primary ciliary dyskinesia epithelial cells *in vitro*. (A) Schematic representation of air-liquid interface culture in which basal cells are cultured to confluence and apical medium removed to stimulate ciliogenesis. The image in the 2D culture is taken from Figure 1B. (B) TEER values in ALI cultures from basal cells expanded in BEGM or 3T3+Y from four healthy and six PCD donors after 28 days of culture. Each point represents a mean value of three consecutive readings from four wells for each individual. *P* values were derived using a two-way ANOVA with Holm-Sidak’s test for multiple comparisons, ns indicates *P* > 0.05. (C-H) qPCR analysis compared the expression of keratin 5 (*KRT5*; C), *P63* (D), *KRT8* (E), *MUC5AC* (F), *MUC5B* (G) and *FOXJ1* (H) between 3T3+Y basal cell cultures and the donor-matched differentiated ALI cultures after 28-30 days of culture. *P* values were derived using two-tailed paired t-tests, ns indicates *P* > 0.05, ** *P* < 0.01 and *** *P* < 0.001. (I) Ciliary beat frequency analysis in ALI cultures from basal cells expanded in BEGM or 3T3+Y from three healthy and seven PCD donors for 28-35 days of ALI culture. *P* values were derived using a two-way ANOVA with Holm-Sidak’s test for multiple comparisons, ns indicates *P* > 0.05.

### PCD phenotypes are preserved in air-liquid interface cultures

A prerequisite for using cultured primary epithelial cells for diagnostic and basic research purposes is that ciliated cells generated in ALI cultures maintain their PCD phenotypes. Cilia were readily detectible using high-speed video microscopy in all cultures, and we found that ciliary beat frequency in ALI cultures from healthy control and PCD patients was consistent between basal cell expansion conditions (Figure 2I). Cilia generated by healthy donor nasal epithelial cells retained a normal ciliary beat pattern in ALI culture (Table 2), as expected based on experiments using human bronchial epithelial cells [8]. Importantly, PCD patient ALI cultures maintained their ciliary beat defects that were present in the original nasal brushings, after basal cell culture and differentiation in both BEGM and 3T3+Y culture conditions. PCD patient cells showed completely static, stiff or dyskinetic cilia that were consistent with the original nasal brushings and that reflected the patterns expected for the respective underlying genetic defects. This was observed from measuring the beat frequency (Figure 2I) and beat pattern, which were very similar between the direct patient observations and the cultured cells (Table 2). The expected ultrastructural defects were also recapitulated (Figure 3). In these experiments, 3T3+Y culture allowed us to analyze ciliary beat frequency and pattern for one healthy donor and three PCD donors for whom failed cultures would have prevented this analysis using traditional BEGM culture (Figure 2I; Table 2).

**Figure 3:**
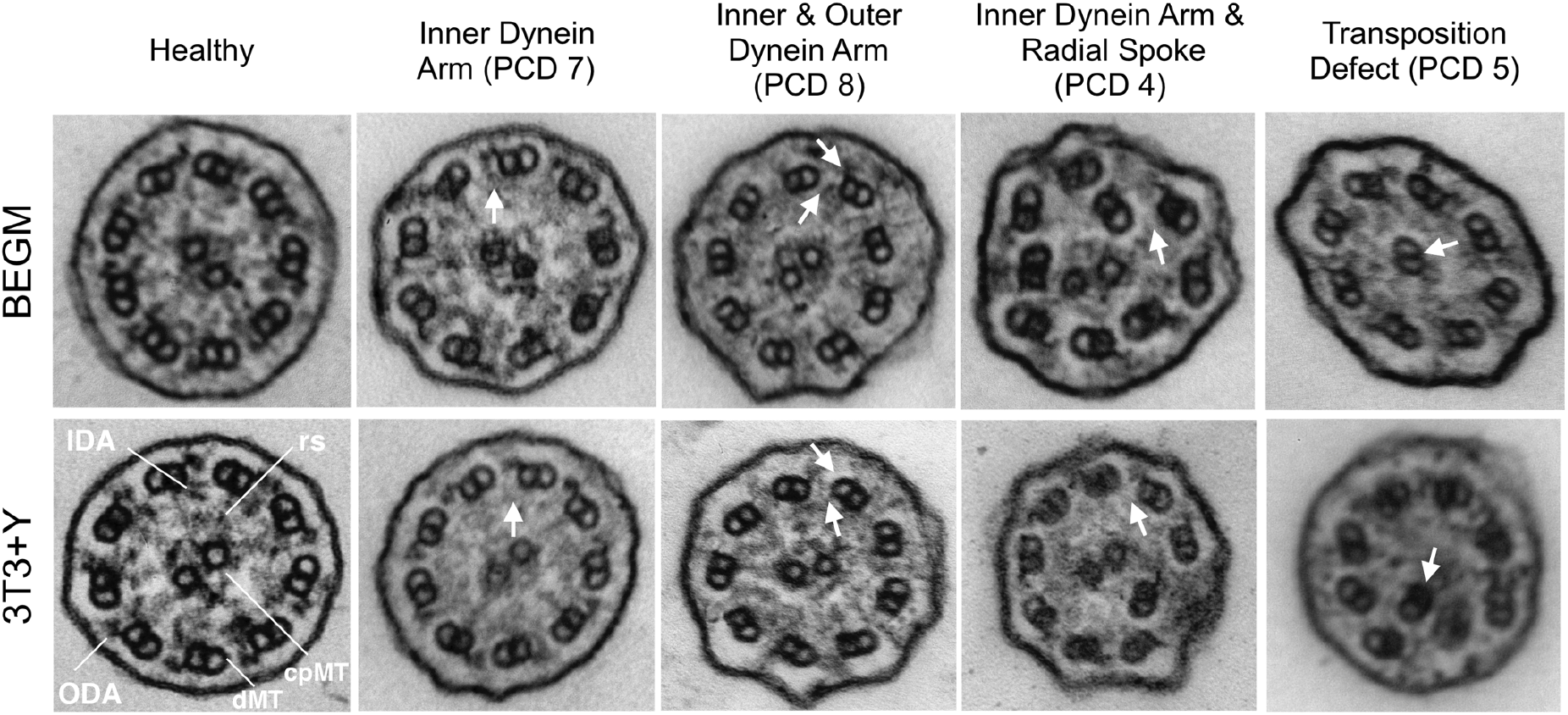
Ultrastructural defects in primary ciliary dyskinesia cell cultures. Electron micrographs of representative ciliary cross-sections illustrating examples of the defects observed in PCD cell cultures. A healthy donor electron micrograph is shown for comparison. The BEGM and 3T3+Y cultures shown are donor-matched. White arrows indicate specific defects and letters indicate: cpMT = central pair microtubules, dMT = double microtubules, IDA = inner dynein arm, ODA = outer dynein arm, rs = radial spoke.

### Miniaturization of PCD air-liquid interface cultures

The ability to expand patient-specific primary PCD epithelial cells with high success rates and assess their ciliary function in culture represents an advance over traditional culture methods for human nasal epithelial cells. However, existing ALI methods are low-throughput. In the knowledge that we can provide large numbers of basal cells as the input from individual patients, we next sought to miniaturize PCD ALI cultures to a 96-transwell format that would be compatible with compound screening using high-content microscopy (Figure 4A). We assessed the ability of cultures from five donors to differentiate in this format, comprising two healthy donors (Healthy 3 and 4, Table 2), two PCD patients with a circular beat pattern (PCD5 and 12, Table 2) and one PCD patient with static cilia (PCD1, Table 2). All donor cultures formed a confluent epithelium containing both MUC5AC^+^ mucosecretory cells and β-tubulin^+^ ciliated cells (Figure 4B), which could be readily visualized using high-speed video microscopy (Supplementary Video File). Analysis of ciliary beat frequency (Figure 4C) and pattern (Figure 4D) showed consistent findings between the 96- and the previously described 24-well format. While a small, statistically significant increase in beat frequency was seen in one healthy donor in the 96-well format (Figure 4C), this is unlikely to be biologically significant as all data points remained within the expected normal range.

**Figure 4:**
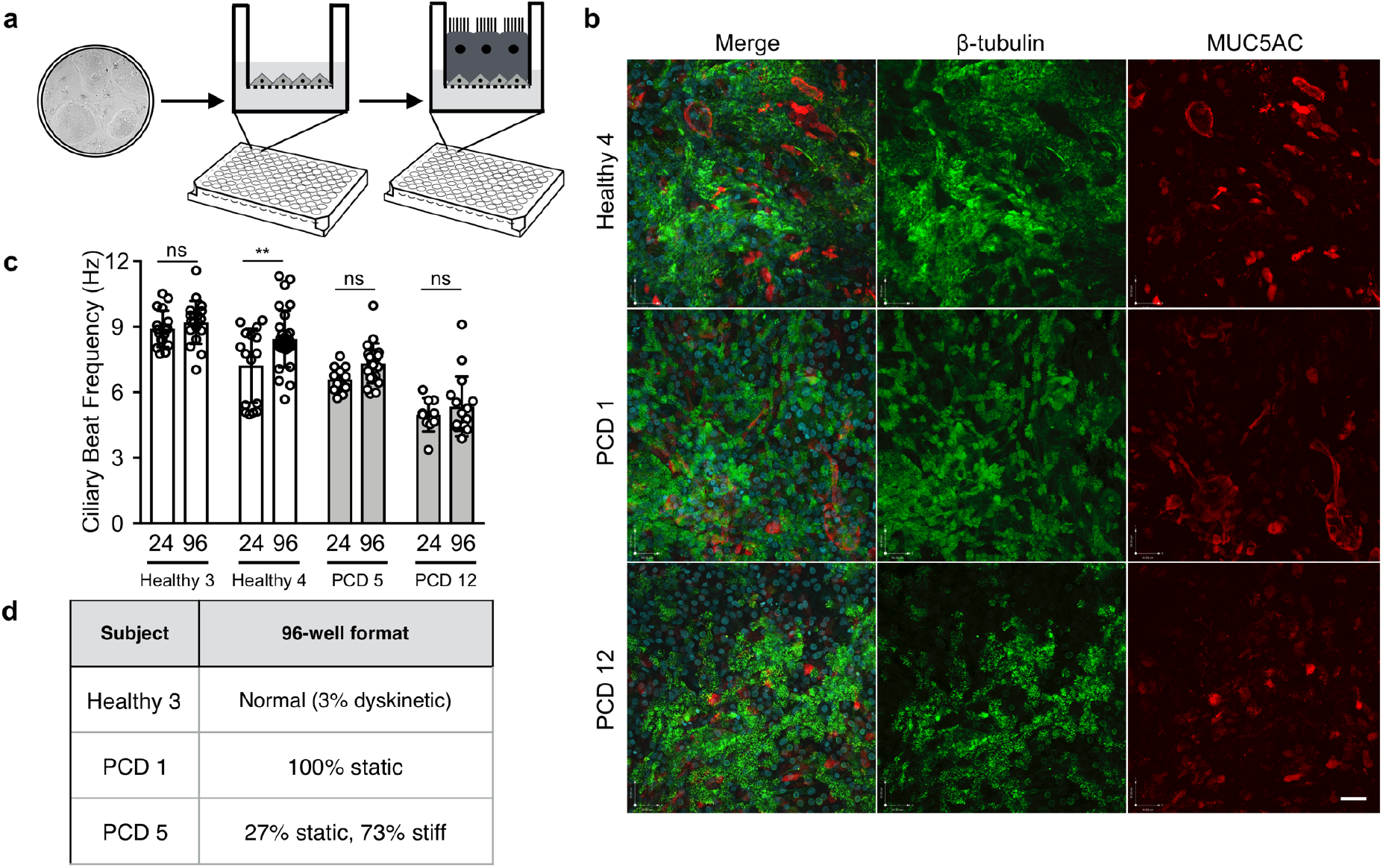
Miniaturization of air-liquid interface cultures to 96-well format to generate PCD patient models. (A) Schematic showing experimental set up for high-content screening in 96-well transwell ALI system. The image in the 2D culture is taken from Figure 1B. (B) Immunofluorescence images demonstrating the presence of β-tubulin-expressing cilia (green) and MUC5AC-expressing mucosecretory cells (red) in 96-well format ALI cultures. Two PCD donors, one with static cilia (PCD 1) and one with circular cilia (PCD 5) defects, are shown in comparison to a healthy donor control. There images are representative of two healthy donors and four PCD donors tested. Scale bar = 50 µm. (C) Ciliary beat frequency analysis comparing multiple recordings from two healthy donors (Healthy 3 and 4) and two PCD patients (PCD 5 and 12) in 24-well and 96-well formats. Data points are individual recordings from 4-5 wells, with 5 videos recorded per well. *P* values were derived using a two-way ANOVA with Holm-Sidak’s test for multiple comparisons, ns indicates *P* > 0.05 and ** indicates *P* < 0.01. (D) Ciliary beat pattern analysis in 96-well format. 24-well equivalent values are found in Table 2.

### Readthrough agents in combination with NMD inhibitors increase basal body formation

Since gentamycin is toxic in humans at serum concentration ranges that achieve readthrough in cell cultures [25] and long-term use of nebulized gentamycin may increase systemic toxicity, we assessed readthrough drugs (gentamicin and ataluren) in combination with nonsense-mediated decay (NMD) inhibitors (amlexanox, escin) in an attempt to improve the therapeutic relevance of our findings. In 96-well format, we treated nasal epithelial cells from patient with a reduced generation of motile cilia (RGMC) ciliopathy, which is caused by a homozygous nonsense mutation (c.441C>A; p.Cys147*) in the *MCIDAS* gene, with dose ranges of gentamicin (50-100 μg/ml) and ataluren (2.5-10 μg/ml) in combination with NMD inhibitors (Figure 5A). Drugs were included from generation of an air-liquid interface onwards, since multicillin expression precedes ciliogenesis [20] (Supplementary Figure 1). After 12 days, we assessed expression of C-Nap1, a centriolar protein that co-localises with β-tubulin^+^ cilia in cultured nasal epithelia [26, 27], using an Opera Phenix high-content screening confocal microscopy system. C-Nap1 expression in healthy control cells showed co-localisation with β-tubulin at day 12 post-ALI (Supplementary Figure 2). In control (untreated) conditions, the patient’s epithelium contained few C-Nap1^+^ basal bodies (Figure 5B, C and Supplementary Figure 3). However, increased basal body formation was seen across various combinations of drug treatments (Figure 5C, Supplementary Figure 3). This is clearly evident with individual C-Nap1^+^ cells visible when analysed by higher resolution confocal microscopy (Figure 5B). *MCIDAS* mRNA was undetectable by qPCR in untreated cultures, but was restored to varying extents after drug treatment was repeated in independent 12-well format ALI cultures (Figure 5D).

**Figure 5:**
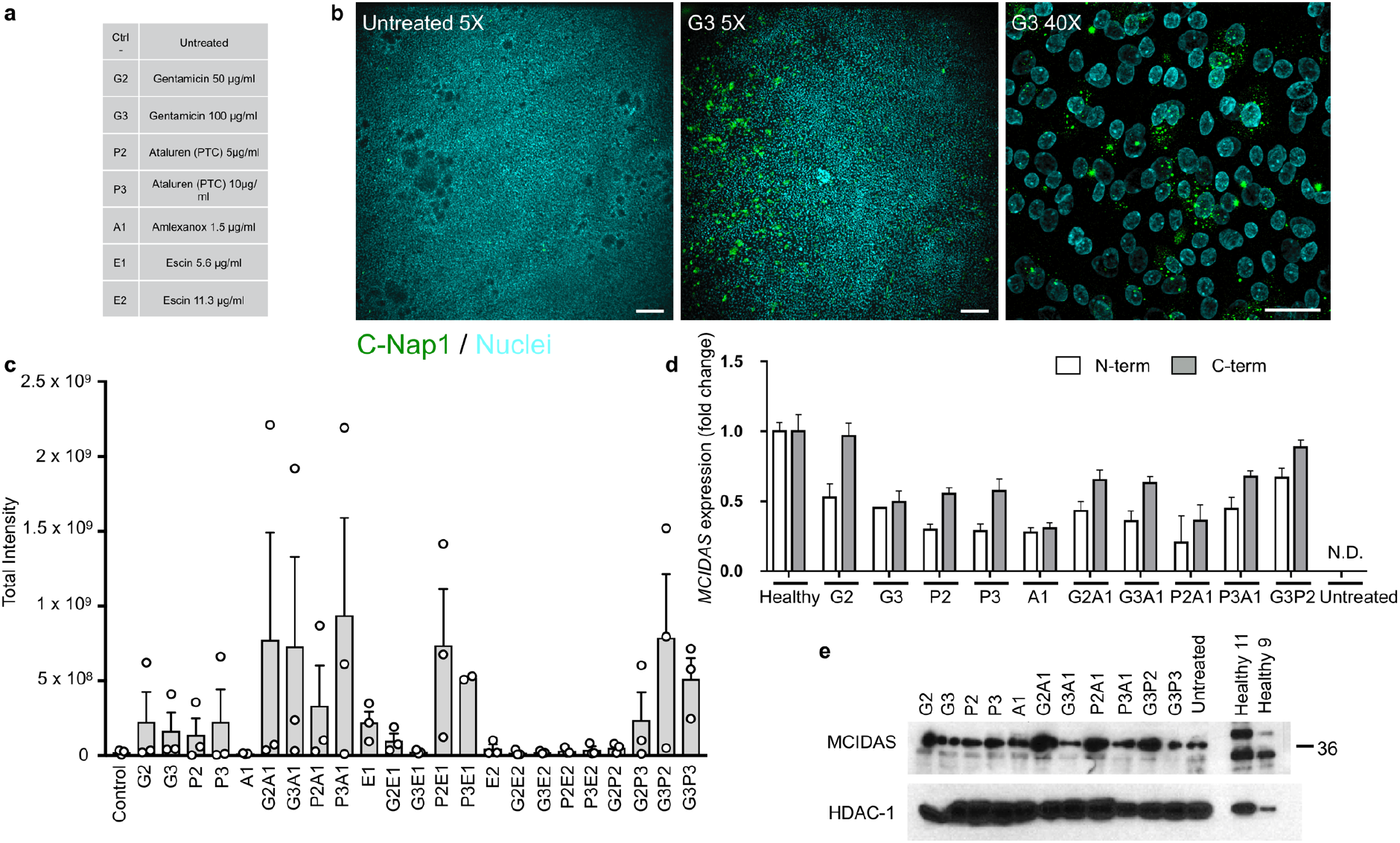
High-content screening to assess readthrough therapies in a patient with *MCIDAS*-mutated RGMC ciliopathy. (A) Key for drug names and doses for high-content screening in patient RGMC 1. (B) An example of basal body formation in the presence of 100 µg/ml gentamicin (G3) readthrough drug treatment shows the high-content immunofluorescence approach based on co-staining of C-Nap1 (a basal body-associated protein; green, FITC) and Hoechst 33258 counterstain (cyan, DAPI). Scale bars = 200 μm (5X images) and 34 μm (40X image). Examples from each condition are shown in Supplementary Figure 3. (C) Quantification the formation of basal bodies after treatment with the combinations of readthrough drugs and nonsense mediate decay inhibitors shown in (A), based on total intensity of C-Nap1 staining, quantified using a custom-made ImageJ macro. An untreated patient control was compared to treated patient samples. No significant differences were found in a one-way ANOVA with Holm-Sidak’s test for multiple comparisons. (D) qPCR analysis comparing *MCIDAS* mRNA abundance between drug-treated conditions on the patient cells versus a healthy control donor in independent cultures using conventional 12-well transwell supports. No mRNA expression was detected in untreated cultures (N.D. = not detected). (E) Western blot analysis of multicillin expression in drug-treated conditions in independent cultures using conventional 12-well transwell supports at day 7 post-ALI.

Recovery of full length multicillin protein was assessed by Western blot in these cultures; increased expression of a band at the expected 38 kDa size was detected in multiple drug treatment conditions, compared to control untreated patient cell cultures and heathy control cultures (Figure 5E). Transmission electron microscopy showed the formation of basal bodies of a range of maturities (Figure 6 and Supplementary Figure 5). Basal body biogenesis is driven by different mechanisms with the canonical mechanism involving duplication of pre-existing centrioles. However, formation of multiple basal bodies in multiciliated cells also follows a non-canonical pathway that is mediated by the formation of structures called deuterosomes [28]. Just prior to deuterosome formation, electron-dense “fibrogranular material” that is enriched in microtubules may be observed [28]. We detected this type of microtubular agglomeration in RGMC cells treated with readthrough agents (Figures 6). These electron dense granules then condense to form hollow, spherical structures called deuterosomes [28], which are also highly electron-dense. Immature centrioles are amplified from daughter centrioles through deuterosome formation [29, 30]. We observed a number of what we believe are deuterosomes in treated RGMC cells (Figure 6 and Supplementary Figure 5). Despite the presence of deutorosomes, we did not observe procentrioles originating from them, although some structured centrioles could be found close to deuterosomes. These centrioles have the characteristic “cartwheel” structure of immature centrioles and are approximately 0.2 μm in diameter, as reported previously [31] (Figure 6).

**Figure 6:**
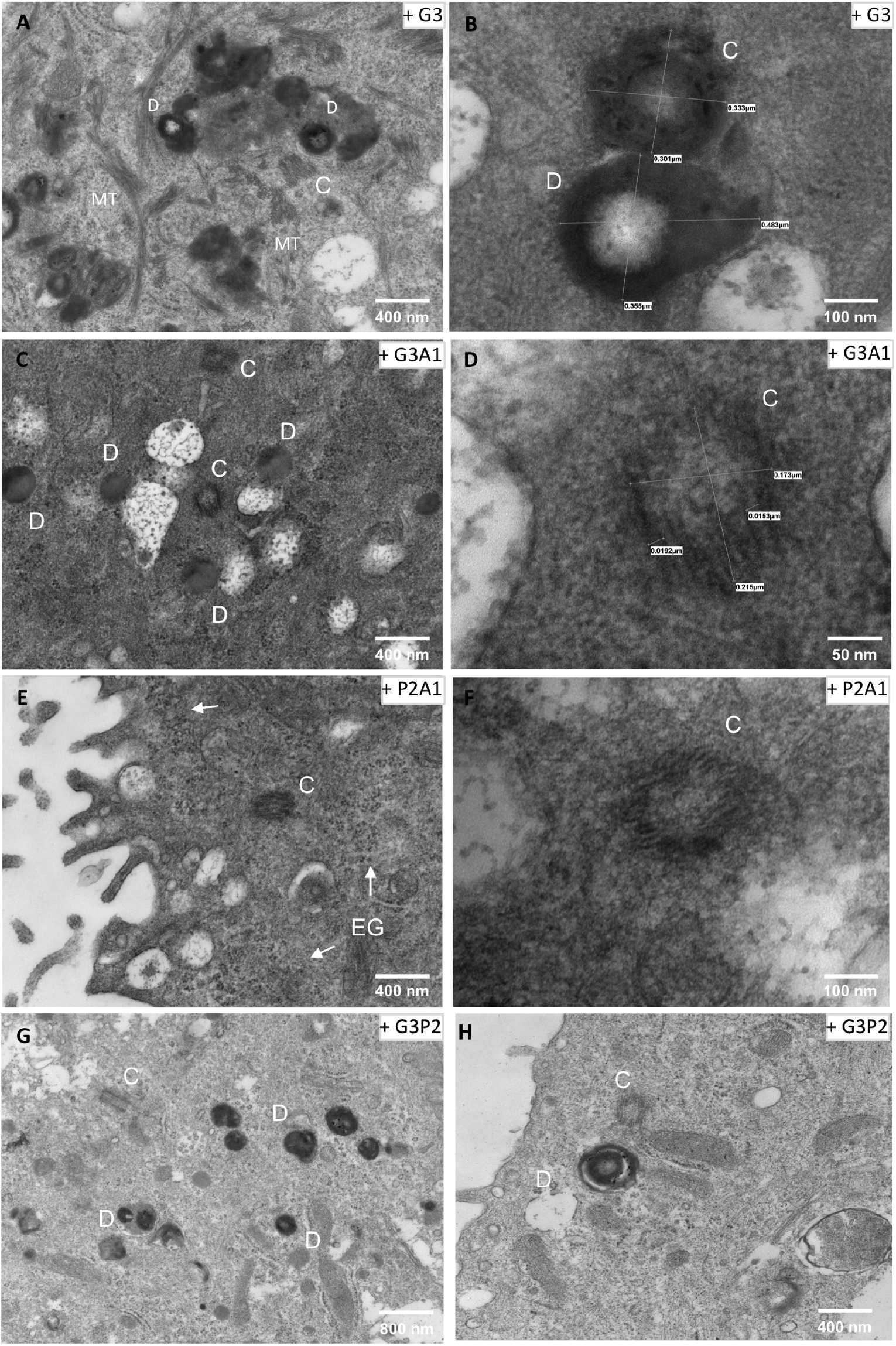
High magnification images of the different structures found in RGMC treated cells. The different panels show representative high magnification images of accumulation of basal bodies “precursors” observed in RGMC cells after treatment with various drugs combinations (as indicated in the top-right corner). The different structures are labelled as: D = deuterosomes, C = centrioles, MT = microtubules agglomeration, EG = electron-dense granules. Electron-dense deuterosomes were observed in high number and we observed several centrioles showing a “cartwheel” structure. Centriole size was ~0.2 μm diameter (Panel C-F, Dawe et al, 2007). None of these structures were observed in untreated controls.

## Discussion

Here we show that, in conditions that allow robust epithelial cell expansion, PCD nasal epithelial cells express proteins characteristic of basal stem/progenitor cells, form differentiated mucociliary epithelia in air-liquid interface cultures and retain patient-specific ciliary dyskinesia phenotypes as previously described for low passage cultures in BEGM [22, 23]. The consistent culture initiation, differentiation capability after extensive culture, absence of viral immortalization [32], and reliable cryopreservation [33] afforded by this approach are a major benefit for PCD research. This will enable the development of large biobanks of well-phenotyped patient-specific PCD epithelial cells of known genotype and repeat experimentation in the same genetic background.

Current therapeutic avenues in PCD focus upon early diagnosis and clinical management in order to prevent lung function decline. However, at present, no therapies that restore ciliary function are available. In the future, the ability to expand significant numbers of human epithelial cells may also open the door to gene editing, mRNA, oligonucleotide, and/or cell therapy approaches in PCD. Notably, cell therapies using epithelial cells expanded on 3T3-J2 feeder layers are approved for clinical use for severe burns and limbal stem cell deficiency [34–36] and expansion from single cell-derived clones is achievable [37]. However, as cellular therapy in genetic lung disease remains a distant prospect [38], model systems that enable the testing of gene therapies or small molecule approaches are much needed. To this end, the production of many cells from individual donors allowed us to miniaturize ALI cultures to 96-well format. In turn, this allowed personalized investigation using high-content, immunofluorescence-based compound screening in the context of a case of RGMC. In a patient with a *MCIDAS* mutation, we demonstrated by immunofluorescence screening of the basal body protein C-Nap1 that combined readthrough and nonsense mediated decay inhibition enhanced basal body formation, although the cultures did not produce motile cilia. This might be due to the limited range of concentrations that we can test in differentiated cultures: at higher doses, compounds like gentamicin, Amlexanox and Escin are toxic. We further demonstrate the feasibility of screening 96-well format cultures using high-speed video microscopy, meaning that this approach could also be taken to screen for agents which correct ciliary beat pattern defects seen in other PCD phenotypes.

Cultures containing only basal epithelial cells are limited for respiratory disease modelling due to the lack of barrier function, the absence of differentiated cell types [39] and the preferential replication of some viruses in specific epithelial cell types (e.g. RSV in multiciliated cells [40]). ALI culture is typically low-throughput and thus not well suited to screening applications. Although drugs were applied basolaterally in cell culture medium, it may be possible to develop a system in which aerosolized drugs are applied to cultures. As such, the approach described here is likely to be widely applicable in basic lung biology, in inhalation toxicology [41] and for investigations of other respiratory diseases with epithelial involvement, such as chronic obstructive pulmonary disease (COPD), asthma and pulmonary fibrosis. While organoid culture has been developed as a scalable primary cell culture model in the context of cystic fibrosis [42], where organoid swelling can be used as a proxy for ion channel function [43, 44], this approach – even with the development of airway organoids [45, 46] – lends itself less well to ciliary diseases, where access to and reliable measurement from the apical surface of the epithelium is essential.

Our approach enables the generation and long-term maintenance of differentiation-competent primary epithelial cell cultures from PCD patients with diverse causative mutations. As a result, it is relevant for the diagnosis of PCD, for basic PCD research and to our knowledge is the first example of a personalized high-content screening approach in a rare respiratory disease using differentiated human respiratory epithelial cells. The culture method and assay miniaturisation allowed identification of a combination of readthrough drugs and NMD inhibitors as an avenue for further pre-clinical exploration in the subset of PCD patients with nonsense mutations that would not have been possible using conventional methods.

## Abbreviations

PCD: Primary ciliary dyskinesia
RGMC: reduced generation of motile cilia
ALI: air-liquid interface
BEBM: bronchial epithelial basal cell medium
BEGM: bronchial epithelial cell growth medium
ROCK: Rho-associated protein kinase
CBF: ciliary beat frequency
NMD: nonsense-mediated decay
IDA: inner dynein arm
ODA: outer dynein arm
MTD-L-IDA: microtubules disorganisation with lack of dynein arm

## Acknowledgements

We are grateful to the families who participated in this study and acknowledge the support of the UK PCD Family Support Group and the COST Action BEAT-PCD (BM1407). We thank Simon Broad and Prof. Fiona Watt (Kings College London, U.K.) for providing the 3T3-J2 fibroblasts that were used in our study.

This work was supported by Action Medical Research (2336: “Restoring the function of cilia in children with primary ciliary dyskinesia - overcoming the genetic block”), by Great Ormond Street Hospital Charity and Sparks National Funding Call 2017/2018 (V4818 - Restoring ciliary function in primary cilia dyskinesia) and by the NIHR Great Ormond Street Hospital Biomedical Research Centre. Microscopy was performed at the Light Microscopy Core Facility, UCL GOS Institute of Child Health supported by the NIHR GOSH BRC award 17DD08. The views expressed are those of the authors and not necessarily those of the NHS, the NIHR or the Department of Health. M.R.F. was funded by the British Council Newton-Mosharafa Fund and the Ministry of Higher Education in Egypt. S.M.J. is a Wellcome Trust Senior Fellow in Clinical Science (WT107963AIA) and receives funding as a member of the UK Regenerative Medicine Platform (UKRMP2) Engineered Cell Environment Hub (MRC; MR/R015635/1) and the Longfonds BREATH lung regeneration consortium. S.M.J. is further supported by The Rosetrees Trust, The Roy Castle Lung Cancer Foundation and The UCLH Charitable Foundation. R.E.H. is a Wellcome Trust Sir Henry Wellcome Fellow (WT209199/Z/17/Z) and is supported by the Roy Castle Lung Cancer Foundation. This work was also supported by the Medical Research Council Core funding to the MRC LMCB (MC_U12266B) and Dementia Platform UK (MR/M02492X/1).

## Materials and methods

### Subjects and genetic analysis

Affected individuals were diagnosed with PCD using standard diagnostic screening by the national commissioning group (NCG)-funded PCD diagnostic service, according to European Respiratory Society diagnostic guidelines [5]. Samples for genetic screening were collected under ethical approval obtained through the London Bloomsbury Research Ethics Committee (08/H0713/82). Informed written consent was obtained from all participants or their guardians prior to enrolment in the study. Genetic screening was performed as previously described, using a targeted next generation sequencing panel approach [47] or as previously reported [48].

### Isolation of epithelial cells from nasal brush biopsies

Human nasal epithelial cell cultures were derived from nasal brush biopsies taken with informed consent. Ethical approval was obtained through the National Research Ethics Committee (REC reference 14/NW/0128) and UCL Research Ethics (reference 4735/001). Biopsies were received on ice in transport medium which consisted of medium 199 (Life Technologies; #22340020) containing 100 U/ml penicillin, 100 µg/ml streptomycin (Gibco; #15290026), 25 µg/ml amphotericin B and 20.5 µg/ml sodium deoxycholate (Gibco; #15290018). Cells were plated directly into T25 flasks in bronchial epithelial growth medium (BEGM; Lonza) and cultured for 7-10 days prior to first passage. After the first passage, cells were either expanded in BEGM or in co-culture with 3T3-J2 fibroblasts as described below.

### Feeder cell culture

3T3-J2 mouse embryonic fibroblasts were grown in Dulbecco’s modified Eagle’s medium (DMEM; Gibco #41966) supplemented with 1X penicillin/streptomycin (Gibco; #15070) and 8% bovine serum (Gibco; #26170). Fibroblasts were cultured at 37**°**C with 5% CO_2_ and medium was changed three times per week. To generate feeder layers, cells were mitotically inactivated by treatment with 4 µg/ml mitomycin C (Sigma; M4287) in culture medium for 2-3 hours. Inactivated fibroblasts were then trypsinized and plated at a density of 20,000 cells/cm^2^ in fibroblast growth medium and allowed to adhere overnight.

### 3T3-J2 co-culture of human nasal epithelial cells

Co-culture with 3T3-J2 fibroblasts was performed as previously described [49]. Epithelial cell culture medium consisted of DMEM (Gibco; 41966) and F12 (Gibco; 21765) in a 3:1 ratio with 1X penicillin/streptomycin (Gibco; 15070) and 5% FBS (Gibco; 10270) supplemented with 5 μM Y-27632 (Cambridge Bioscience; Y1000), 25 ng/ml hydrocortisone (Sigma; H0888), 0.125 ng/ml EGF (Sino Biological; 10605), 5 μg/ml insulin (Sigma; I6634), 0.1 nM cholera toxin (Sigma; C8052), 250 ng/ml amphotericin B (Fisher Scientific; 10746254) and 10 μg/ml gentamycin (Gibco; 15710). Streptomycin and gentamycin were removed from the medium for RGMC cell culture. Epithelial cells were cultured at 37**°**C and 5% CO_2_ with three changes of medium per week. Population doublings (PD) were calculated as PD = 3.32 * (log (cells harvested / cells seeded), 10). Cryopreservation was performed using Profreeze freezing medium (Lonza) according to the manufacturer’s instructions and using epithelial cell culture medium as defined above to dilute the stock solution.

### Differentiation in air-liquid interface cultures

At confluence, basal cells were separated from feeder cells using differential trypsinization [8], where a brief initial incubation with 0.05% trypsin/EDTA was sufficient to remove fibroblasts but epithelial cells remained adherent. After washing, a second, longer incubation efficiently removed epithelial cells. Basal cells were seeded on collagen I-coated, semi-permeable membrane supports (Transwell-Col, 0.4 µm pore size; Corning) in submerged culture in BEGM containing 5 µM Y-27632. For 12-well transwells, 1 × 10^6^ cells were seeded per membrane in 250 µl medium. Miniaturised ALI cultures were performed in 96-transwell plates (Corning; 3380; 1 µm pores, polyester membrane) and 0.2 × 10^6^ cells were seeded per membrane in 75 µl medium. Upon confluence, cells were fed only from the basolateral side with air-liquid interface medium. Air-liquid interface medium consisted of 50% BEGM and 50% hi-glucose minimal essential medium (DMEM, Gibco) containing 100 nM additional retinoic acid (Sigma Aldrich). For 96 transwell ALI cultures, BEGM was replaced with Promocell’s Airway Epithelial Cell Growth Medium. Medium was exchanged three times per week and gentle washing with BEBM or Promocell basal medium removed the mucus produced on the apical surface once per week.

### Immunofluorescence

Cultured basal cells were seeded in 8-well chamber slides or fixed directly on the transwell membrane by incubation in 4% paraformaldehyde for 30 minutes at room temperature. Cells were stored at 4**°**C in PBS until the time of staining. Cells were blocked and permeabilized using block buffer (3% BSA in PBS containing 0.01% Triton X-100) at room temperature for 1 hour, prior to overnight staining with primary antibody (in 1% BSA in PBS) at 4°C. Primary antibodies used were anti-C-Nap1 (Sigma; WH0009918M1; 1:50) for basal body staining, anti-β-tubulin (Abcam; ab15568; 1:100) and anti-MUC5AC (Invitrogen; 1:100). Cells were washed three times in PBS for 5 minutes and secondary antibody (in 1% BSA in PBS; Molecular Probes; AlexaFluor dyes) was applied for 2 hours at room temperature. Hoechst 33258 staining solution (Sigma) was applied for 20 minutes at room temperature as a nuclear counterstain prior to imaging.

### High-content screening in miniaturised ALI cultures

Cells from the patient whose cells were examined in the high-content screening assays were isolated and expanded in 3T3+Y cell culture conditions as described above. Gentamicin (50 and 100 μg/ml) and ataluren (5 and 10 μg/ml) alone or in combination with amlexanox (1.5 µg/ml) and escin (5.6 and 11.3 µg/ml) were applied to ALI cultures basolaterally from the onset of differentiation (i.e. at air-lift). Drugs were refreshed each time the cultures were fed. Cells were fixed for immunofluorescence at day 12 post-ALI for assessment of basal body formation. Selected drugs and drug/combinations from the screening were also applied to 12-well transwell ALI cultures and cells were collected for western blot and qPCR analysis after 7 days.

For assessment of basal body formation, cells were imaged directly in 96-well transwells using automated confocal microscopy (Opera Phenix High-Content Screening System, PerkinElmer, 5x objective). For higher magnification imaging, cells were mounted in 80% glycerol, 3% n-propylgallate (in PBS) mounting medium and images were obtained using an inverted Zeiss LSM 710 confocal microscope (Zeiss). Analysis of basal bodies was performed by using a custom ImageJ macro; the analysis approach is summarised in Supplementary Figure 3 and the macro is freely available at the following link: https://github.com/DaleMoulding/Fiji-Macros/blob/master/DanielaSpotClustersv9.ijm.

### Transepithelial electrical resistance (TEER)

TEER values were measured using an EVOM2 resistance meter and Endohm chamber (World Precision Instruments) with cup size appropriate for the size of culture insert (6 mm culture cup for 24-well transwells and 12 mm culture cup for 12-well transwells). Transwells were placed into the culture cup and readings were taken after the TEER reading had stabilized (typically 5-10 seconds). Readings were taken from three independent transwells once cultures were fully differentiated, i.e. after at least 5 weeks of ALI culture, to obtain an average TEER value for each culture (9-12 readings per donor).

### Ciliary beat frequency and beat pattern

For ciliary analyses, the ciliated epithelium was removed from the transwell insert by gentle scraping with a spatula. Cells were washed with transport medium (as described above) and dissociated by gentle pipetting. The cell suspension was divided to allow ciliary beat frequency and beat pattern analyses on unfixed cells and TEM analysis on cells fixed in glutaraldehyde.

Beating cilia were recorded using a digital high-speed video camera (Motion Pro 4x; IDT) at a rate of 500 frames/second using a 100x objective [50]. Ten videos were acquired for each donor and CBF of individual ciliated cells was determined by counting the number of frames required for 5 full sweeps of a clearly visible ciliary tip. This was converted to CBF, where CBF = 500 / (number frames for 5 beats) x 5. The dyskinesia index presented is the percentage of dyskinetic ciliated cells relative to the total number of motile ciliated cells [51]. To analyze ciliary activity in cells differentiated in 96-well transwell ALI cultures, nasal epithelial cells were observed using an inverted microscope system (Nikon Ti-U) with a 20x objective. At least 20 top-down videos per donor were recorded and CBF was analyzed using the ImageJ plugin CiliaFA [52].

### Quantitative real-time PCR (qPCR)

Cultured epithelial cells were collected, mRNA was extracted in TRIzol and stored at −80**°**C. Cultures containing 3T3-J2 fibroblasts were differentially trypsinized to remove feeder cells as described above. After thawing on ice, total RNA was isolated using a Direct-zol RNA Isolation Kit (Zymo Research). RNA quantity was assessed using a Nanodrop One system and reverse transcription was performed using qScript cDNA SuperMix (Quantabio). The following Taqman pre-designed, inventoried probes, along with 2x PCR Master Mix (Applied Biosciences), were used for qPCR reactions: *KRT5* (Hs00361185_m1), *KRT14* (Hs00265033_m1), *KRT8* (Hs01595539_g1), *TP63* (Hs00978339_m1), *NGFR* (Hs00609977_m1), *ITGA6* (Hs01041011_m1), *MUC5AC* (Hs01365616_m1), *MUC5B* (Hs00861595_m1), *FOXJ1* (Hs00230964_m1) and *B2M* (Hs00187842_m1). qPCR was performed under standard conditions using an Eppendorf Real-Time PCR machine. Relative RNA quantitation was based on deltaCT calculations and all samples were compared to the housekeeping gene β2-microglobulin. For *MCIDAS* detection, primers (Table 1) and Power SYBR™ Green PCR Master Mix (Thermo Fisher Scientific) were used in standard conditions and relative RNA quantification was normalized to the housekeeping gene *GAPDH*.

**Table 1:**
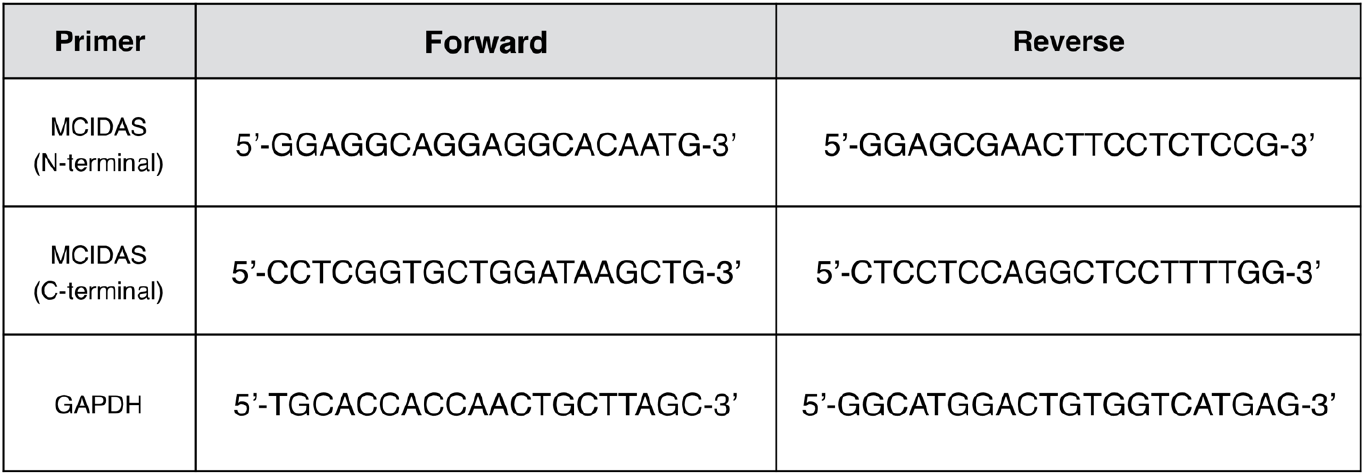
Primers used for *MCIDAS* qPCR.

### Western blot

Cells were collected from transwells by scraping with a spatula, pelleted and washed twice with PBS. The nuclear protein fraction was extracted with NE-PER™ Nuclear and Cytoplasmic Extraction Reagents (Thermo Fisher Scientific) and the concentration determined with Pierce BCA assay kit (Thermo Fisher Scientific). Nuclear extracts were fractionated by Sodium Dodecyl Sulphate-Polyacrylamide Gel Electrophoresis (SDS-PAGE) and transferred to a PVDF membrane (for 1 hour at 100V). Membranes were blocked in 5% (w/v) milk in PBST (3.2 mM Na_2_HPO_4_, 0.5 mM KH_2_PO_4_, 1.3 mM KCl, 135 mM NaCl, 0.05% Tween® 20, pH 7.4) for 1 hour at room temperature and then incubated at 4°C overnight with primary antibodies (anti-multicillin, Sigma Aldrich, HPA058073 1:500; anti-HDAC1, NovusBio, 1:500). Membranes were then washed three times in PBST and incubated with horseradish peroxidase-conjugated anti-rabbit antibodies (Cell Signalling Technologies; 1:3,000) for 1 hour. Blots were washed with PBST three times and developed with an ECL system (Biorad) according to the manufacturer’s protocol.

### Transmission electron microscopy (TEM)

Epithelial cells were scraped from transwells into M199 medium and fixed in 4% glutaraldehyde in Sorensen’s phosphate buffer (pH 7.4). Cells were then re-suspended in Sorensen’s buffer and stored at 4°C until they were further processed through to resin as previously described [51]. Ultrathin (70 nm) sections were cut and collected on copper grids, stained in 1% uranyl acetate, counterstained in Reynold’s lead phosphate and examined by TEM [53].

### Statistics

Statistical analyses were performed using GraphPad Prism using the statistical tests indicated in figure legends.

## Data Supplement

**Supplementary Figure 1.**
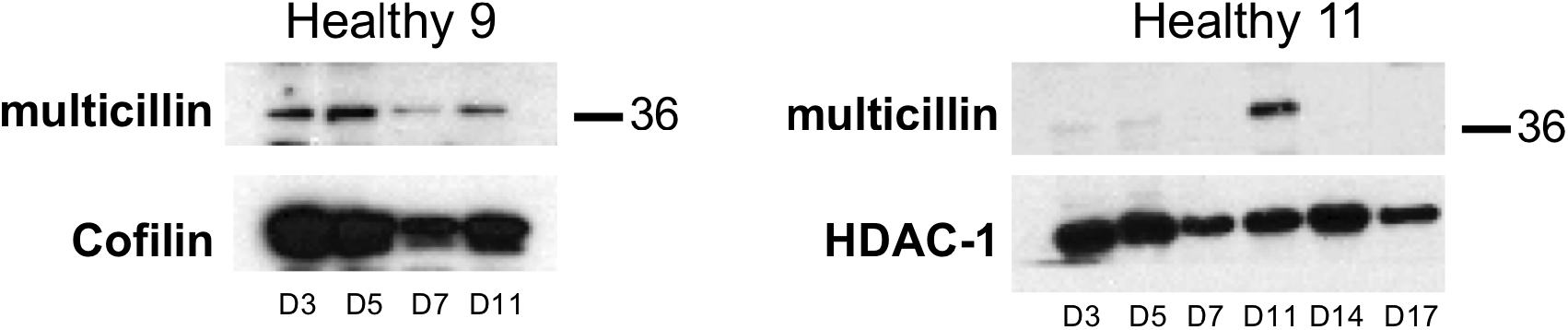
Western blots of multicillin in healthy volunteers at ALI. Cells were collected at different time points (days) after transfer into air-liquid interface culture. SDS page of nuclear extracts from cultures from two different healthy volunteers, showing multicillin expression compared to cofilin or histone deacetylase (HDAC-1).

**Supplementary Figure 2:**
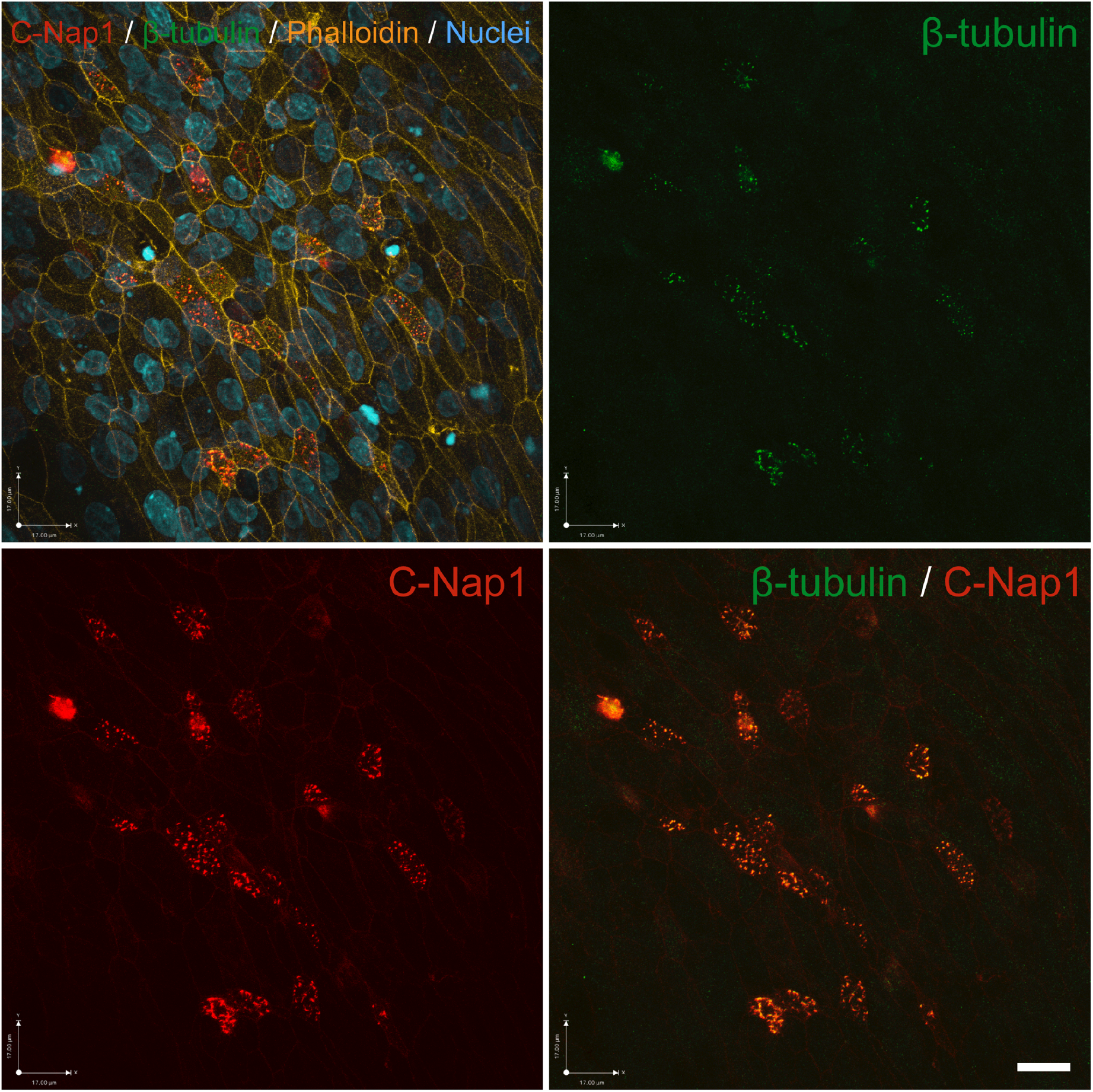
Formation of basal bodies in healthy donor cells in 96 transwell air-liquid interface cultures. Immunofluorescence images demonstrating the colocalization of C-Nap1 (red) basal body marker with the β-tubulin (green) cilia marker in healthy cells grown at ALI in 96 transwell for 15 days. Nuclei are in blue (DAPI) and F-actin in orange (phalloidin). Scale bar = 17 µm.

**Supplementary Figure 3:**
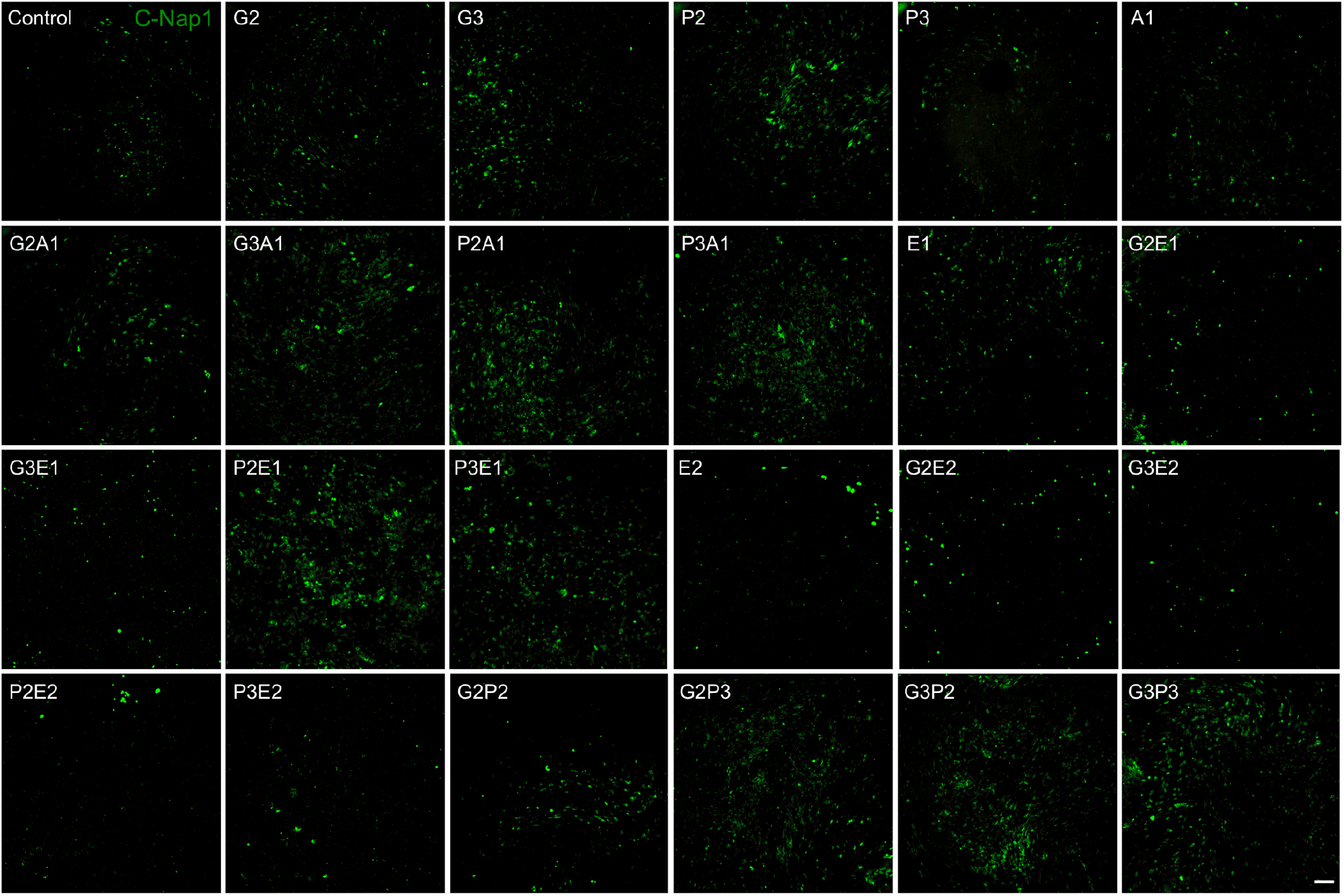
Confocal scanning images from screening of RGMC cells treated with read-through drugs in ALI cultures. C-Nap1 staining (green), Z-stack projection, max intensity. Scale bar = 200 µm.

**Supplementary Figure 4:**
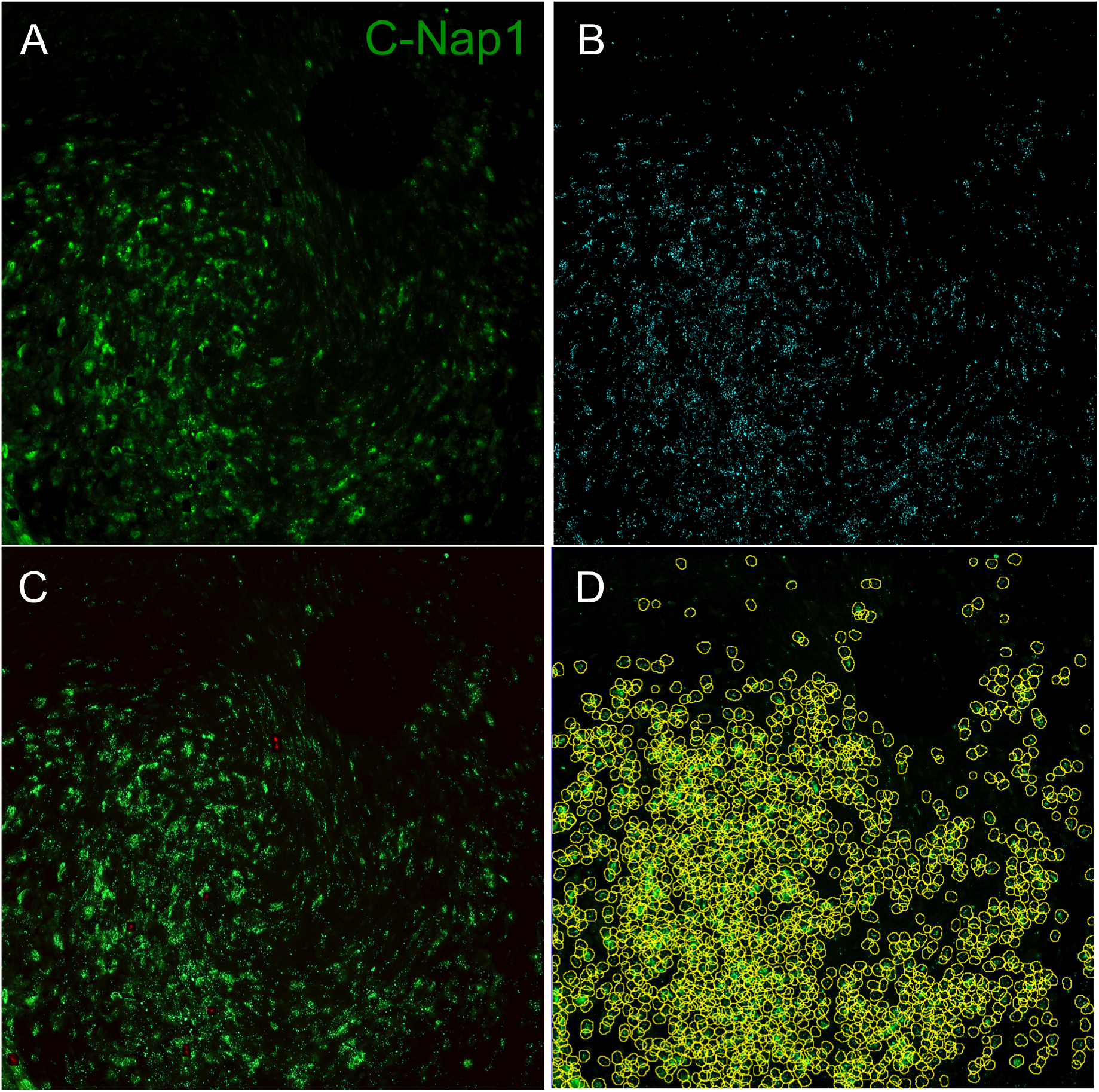
Analysis of basal bodies in cultured primary human nasal epithelial cells at air-liquid interface using ImageJ. Stacks from confocal scanning of 96 transwell plate were analysed in ImageJ. (A) original image (C-Nap1 staining (green), Z-stack projection, max intensity). (B) Selected brightest points, with a radius of 4 pixels. (C) Over-saturated areas where excluded from analysis (red areas). (D) clusters of points (i.e. C-Nap1 staining; minimum of 4 points per cluster at a distance of 15 pixels). Scale bars = 200 µm.

**Supplementary Figure 5:**
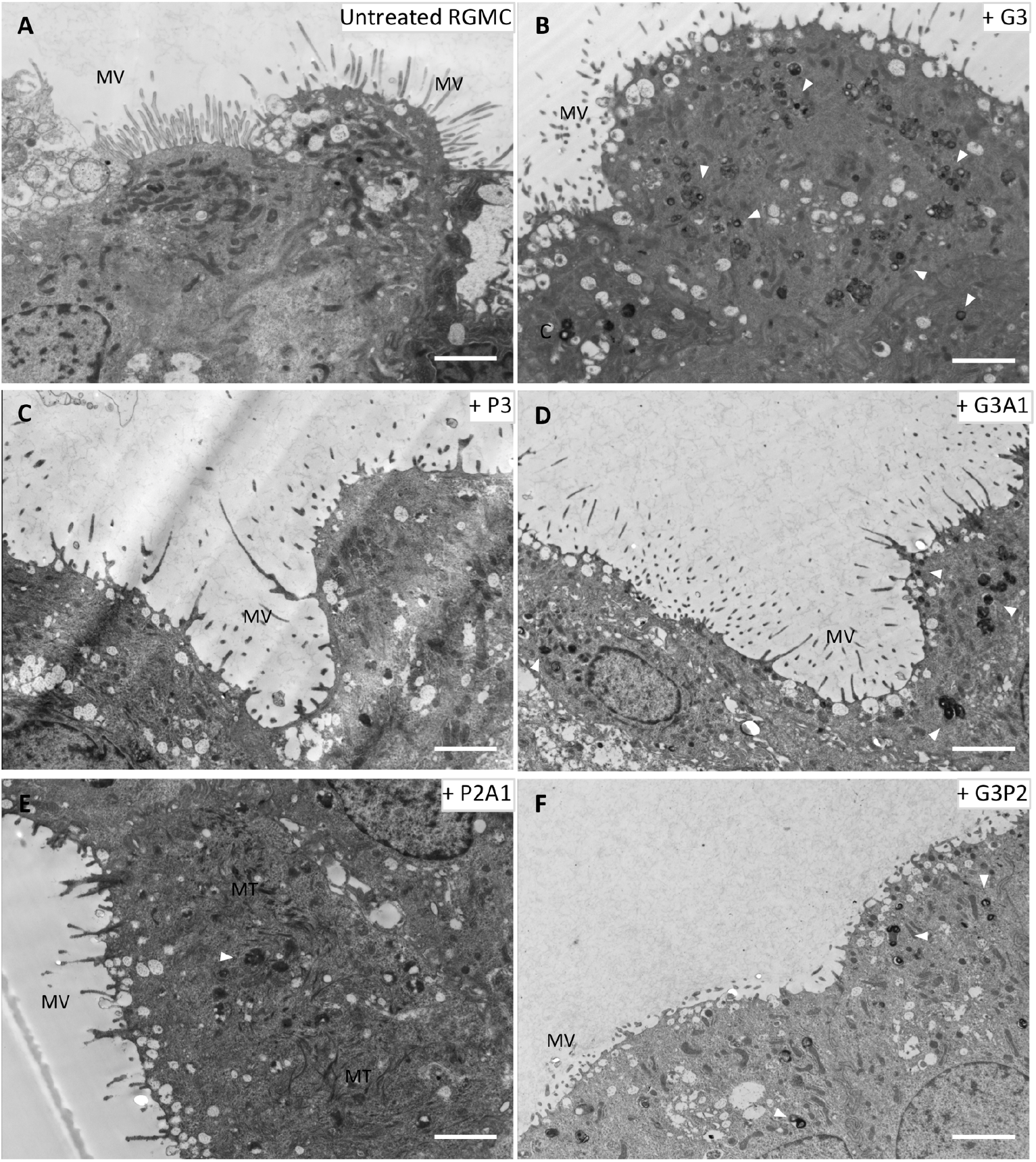
Transmission electron microscopy of basal body precursors following read-through therapy in cells from a patient with *MCIDAS*-mutated RGMC ciliopathy. The different panels show representative low magnification images of RGMC cells untreated (A) and treated with different drugs combinations: Gentamicin 100 μg/ml (B, G3); Ataluren 10 μg/ml (C, P3); Gentamycin 100 μg/ml and Amlexanox 1.5 μg/ml (D, G3A1); Ataluren 5 μg/ml and Amlexanox 1.5 μg/ml (E, P2A1), Gentamycin 100 μg/ml and Ataluren 5 μg/ml (F, G3P2). Cells cultured at ALI were fixed at day 12 after air-lifiting. As seen in the untreated control these cells present long microvilli (MV) and not cilia. After drugs treatment precursors of basal bodies can be identified: electron-dense deuterosomes are indicated with arrowheads, we can distinguish some more structured centrioles (labelled with C) and microtubules agglomerations (MT). Scale bar = 2 μm.

